# *In situ* treatment of H. pylori infection in mice stomach with bioengineered probiotic bacteria releasing guided Antimicrobial peptides

**DOI:** 10.1101/2021.06.11.448139

**Authors:** Ankan Choudhury, Patrick Ortiz, Christopher M. Kearney

## Abstract

**Objectives:** Targeted therapies seek to selectively eliminate a pathogen without disrupting the resident microbial community. This is even more important when a pathogen like *H. pylori* resides in stomach, a sensitive microbial ecosystem. Using a probiotic like *Lactococcus lactis* and bioengineering it to release a guided Antimicrobial Peptide (AMP) targeted towards the pathogen offers a pathway to specifically knock-out the deleterious species and not disturbing the stomach microbiome.

**Results:** Three AMPs, Alyteserin, CRAMP and Laterosporulin, were genetically fused to a guiding peptide MM1, which selectively binds to Vacuolating Toxin A (VacA) of *H. pylori* and cloned into an excretory vector pTKR inside *L. lactis*. The probiotics were then fed to mice infected with *H. pylori*, both as a therapeutic and prophylactic measure, and the samples were collected using a novel gavage method and analyzed using qPCR and Illumina sequencing of the extracted stomach samples over a 10-day period. Microbiome analysis with Next-Gen sequencing also revealed a dysbiosis created by *H. pylori*, determined by creating a Correlation network model with the relative abundances of taxa across the samples, and this dysbiosis was palliated by the bioengineered probiotics which preserved and boosted key microbiome species and reducing the load of deleterious ones. The bioengineered probiotic also significantly improved the OTU diversity compared to antibiotics and *L. lactis* cloned with empty vector, with gAMP-*L. lactis* faring the best.

**Conclusions:** Probiotics bioengineered to excrete guided AMPs can be a novel and useful approach for combating pathogens without endangering the natural microbial flora. Given the wealth of AMPs and guiding ligands, both natural and synthetic, this approach can be adapted to develop a diverse array of chimeric guided AMPs and can be cloned into probiotics to create a safe and effective alternative to conventional chemical antibiotics.

## Introduction

Current treatment regimens for *H. pylori* infections often include triple and quadruple antibiotic therapies to match the growing challenge of antibiotic resistance. Such regimens include combinations of amoxicillin, tetracycline, bismuth, metronidazole, clarithromycin, and more. Alongside such vigorous employment of antibiotics, quadruple, quintuple, and sextuple antibiotic-resistant strains of *H. pylori* have arisen (Boyanova et al., 2019), worsening the predicament. This development of antibiotic resistance in *H. pylori* has made it urgent to develop new therapeutic strategies to combat infection. The broad spectrum antibiotics which function as rRNA inhibitors, β-lactams, nucleic acid inhibitors also deleteriously effect off target bacteria, and a often these antibiotics while trying to curb a single infection increases the dysbiosis of commensal (Becattini et al., 2016; Langdon et al., 2016; Zarrinpar et al., 2018). Antibiotic-associated Dysbiosis often precipitates into intestinal inflammatory diseases like colitis (Strati et al., 2021), worsens neuro-immune mechanisms and viscerosensory functionalities (Aguilera et al., 2015) and often makes way for bloom of pathogens (Vangay et al., 2015) creating other possibly more serious infectious diseases. This keeps exacerbating as we are creating newer, stronger small molecule antibiotics to kill multi drug resistant bacteria with ever-evolving resistance mechanisms, but then these stronger antibiotics also kill a wider variety of commensal bacteria causing dysbiois.(Becattini et al., 2016; Langdon et al., 2016; Zarrinpar et al., 2018).

To develop a therapy strictly active against *H. pylori*, we used antimicrobial peptides (AMP) conjugated with a guiding moiety that will make it attach specifically to the target pathogen. We have demonstrated the efficacy of such method in Chapter Two by using a guiding peptide, discovered in previous literature through biopanning experiments, to modify the bactericidal activity of two AMPs and making them specifically kill *Staphylococcus* bacteria. In the previous chapter we used three AMPs (CRAMP, Laterosporulin and Alyteserin) (Baindara et al., 2016; Conlon et al., 2010, 2009; Hase et al., 2003; Neshani et al., 2019; Singh et al., 2015; Zhang et al., 2016, 2013) that have been reported to have bactericidal action towards Gram negative bacteria and conjugated a fragment of a human platelet protein Multimerin 1 to it. A fragment of Multimerin 1 (AA 321-340) was found to specifically bind to VacA toxin (Satoh et al., 2013), a toxin released by *H pylori* which often remains on the surface of the bacteria (Foegeding et al., 2016; Ilver et al., 2004; Voss et al., 2014). Using this fragment MM1 we have created chimeric gAMPs that were cloned into *L. lactis* using a modified pT1NX vector (pTKR) that releases the expressed AMP/gAMP using a secretory signal usp45 and triggered by a low pH promoter P1(Steidler et al., 2004, 1998; van Asseldonk et al., 1990; Waterfield et al., 1995).

In our previous chapter we demonstrated how the *L. lactis* cloned with AMPs and their corresponding gAMPs showed efficacy in killing *H. pylori*, with the gAMP-cloned *L. lactis* showing a preferential bactericidal activity towards *H. pylori* by showing a diminished activity towards other non-target bacteria like *Lactobacillus plantarum* and *E. coli*. This was also repeated when using chemically synthesized Alyteserin, one of the three AMPs used to clone the bacteria with, and its guided counterpart MM1-Alyteserin. MM1-Alyteserin showed similar MIC towards *H. pylori* as Alyteserin but had significantly higher MIC when used against a non-target bacteria *E. coli*, proving that the guiding moiety made its activity specific towards *H. pylori*. This would be beneficial in killing only the pathogen we desire while not harming the rest of the native microbiome. To determine the efficacy of the therapy *in vivo* we administered C57BL/6J mice with the cloned *L. lactis* that has been infected with *H. pylori*. To check if the effect of the therapy can also be achieved as a prophylactic measure, we also administered the mice with bioengineered *L. lactis* before being given *H. pylori*. The stomach samples of these mice were collected using a novel gavage method and were sequenced using Illumina Miseq with primers targeted towards the 16s V4 region (Caporaso et al., 2012, 2010). Further analysis of the sequencing data was performed with QIIME2 (Bolyen et al., 2019) and CCREPE (Schwager et al., n.d.), for the correlation network analysis on differential taxa abundance. This chapter further explores the Methods undertaken for this research and the observations we made in the end.

## Materials and Methods

### Administering L. lactis and H. pylori in mice by oral gavage and sample collection

The *L. lactis* cultures were propagated overnight GM17 broth with erythromycin (5 μg/ml) and no shaking. The overnight cultures were spun down at 4000g for 15 min at 4°C. The pellets were resuspended in sterile PBS. *H. pylori* SS1 stocks were grown overnight on Blood-TS agar under microaerobic condition and >5% CO2 environment and then scraped by a sterile loop and resuspended in sterile PBS. Both the bacterial suspensions were fed to the mice using 1.5 oral gavage needle not exceeding half their stomach volume (~250 μL). The CFUs of the resuspension being fed were determined by diluting the resuspension 1/1000 and 1/10000 times and plating on appropriate plates. For both *L. lactis* and *H. pylori*, the inoculum sizes were kept ~50,000 CFUs/μl.

Pre and post inoculation samples from the mouse stomach were collected by flushing their stomach with excess PBS (~250 μL). The mice were fed the PBS using a gavage needle that reaches well into the stomach. Then without losing the suction and removing the needle out of the mouse esophagus, the plunger is moved up and down twice without drawing any substantial volume of fluid out. The presence of a negative pressure during pulling the plunger is preferrable. The plunger is then completely pulled out which will draw out around 50-75 μl of stomach fluid. The collected fluid is then put in tubes/vials for storage. The fluid should be slightly viscous and may or may not have suspended food fragment and /or mucus.

4 schemes of bacterial inoculation were followed: *H. pylori* followed by *L. lactis, H. pylori* followed antibiotic (Amoxycillin:Tetracycline :: 4.5:4.5 mg/25 g of mice)*, L. lactis* followed by *H. pylori* and only *H. pylori*.

For scheme 1 (Therapeutic groups): Stomach samples were collected on Day 0 before *H. pylori* inoculation; resuspended *H. pylori* were fed by oral gavage once daily for 3 consecutive days; stomach samples were then collected on Day 5 to test for *H. pylori* presence and on Day 5 resuspended *L. lactis* (cloned with either empty vector, AMP or gAMP) were fed to the mice; subsequent stomach samples were collected on Day 8 and 10.

For scheme 2 (Antibiotic group): Stomach samples were collected on Day 0 before *H. pylori* inoculation; resuspended *H. pylori* were fed by oral gavage once daily for 3 consecutive days; stomach samples were then collected on Day 5 to test for *H. pylori* presence and on Day 5 antibiotic cocktail was fed to the mice; subsequent stomach samples were collected on Day 8 and 10.

For scheme 3 (Prophylactic groups): Stomach samples were collected on Day 0 before *L. Lactis* inoculation; stomach samples were taken on Day 3 followed immediately by inoculation with *H. pylori* by oral gavage once daily for 3 consecutive days; stomach samples were then collected on Day 8 and 10 to test for *H. pylori* presence.

For scheme 4 (Null Control group): Stomach samples were collected on Day 0 before *H. pylori* inoculation; resuspended *H. pylori* were fed by oral gavage once daily for 3 consecutive days; stomach samples were then collected on Day 5, 8 and 10 to test for *H. pylori* presence.

6 mice were used per AMP and gAMP both for therapeutic (scheme 1) and prophylactic (scheme 3). 6 mice were also used to create the null control group (scheme 4), antibiotic treatment group (scheme 2) and the empty vector (pTKR) group for both therapeutic and prophylactic schemes.

### PCR determination of presence of bioengineered L. lactis in mice stomach

The stomach samples of mice at Day 10, 5 days after feeding them cloned *L. lactis*, were subjected to a PCR test (NEB Taq Polymerase, 95°C Denaturation for 5 minutes; 30 cycles of 95°C Denaturation for 30 s, 60°C Annealing for 15 s, 68°C Extension for 30 s; Final extension for 2 minutes) using primers specific to the pTKR vector (forward: 5’ – GCCTGAGCGAGACGAAATAC – 3’, reverse: 5’ – TTATGCCTCTTCCGACCATC – 3’). The PCR products were ran on a 1% agarose gel to ascertain the size of the products.

### Assay for determining H. pylori amount in mice stomach by qPCR

The stomach samples collected were heated at 100°C for 15 min and chilled at 4°C for 5 min. The supernatants were collected and plated in 96 well plate and qPCR was performed with primers for VacA gene (forward: 5’-ATGGAAATACAACAAACACAC-3’, reverse: 5’-CTGCTTGAATGCGCCAAAC-3’) to quantify for *H*. pylori. Standard curves for *H. pylori* against the C_T_ values were constructed by including different dilutions of the overnight cultures of *H. pylori* (1/10, 1/100, 1/1000, 1/10000) in the qPCR and plating those dilutions on respective plates to determine the corresponding CFU/μl values. The CFU/ μl values for each sample were determined by plotting the C_t_ values against the standard curve previously built.

### Analysis of mice stomach microbiome affected by L. lactis and H. pylori inoculation

The stomach samples collected were heated at 100°C for 15 min and chilled at 4°C for 5 min. The supernatants were collected and plated in 96 well plate for upstream processing for Next Gen sequencing. The samples were amplified with 16s V4 primers (forward: 5’-TCG TCG GCA GCG TCA GAT GTG TAT AAG AGA CAG GTG YCAGCMGCCGCGGTAA-3’, reverse:5’-GTC TCG TGG GCT CGG AGA TGT GTA TAA GAG ACA GCC GYCAATTYMTTTRAGTTT-3’) (Caporaso et al., 2012, 2010) and then with Illumina index primers (“Indexed Sequencing Overview Guide (15057455),” n.d.) with subsequent clean-up and purification. The samples were pooled into a library, spiked with PhiX DNA and sequenced using Illumina Miseq v3 kit. The data was demultiplexed, denoised and analyzed for taxonomic abundance using QIIME2 (Bolyen et al., 2019). The alpha and beta diversity analysis were also performed using QIIME2. The taxonomic abundance data (at Genus level) was analyzed using the CCREPE package in R (Schwager et al., n.d.) with the microbial community of the mice stomach at Day 0 compared against the community from the samples taken at Day 5 after 3 consecutive days of *H. pylori* inoculation, to determine the correlation between the taxa which went up or down in relative abundance on addition of *H. pylori*.

## Results

### L. lactis cloned with pTKR were still present in the mice stomach after 5 days

The samples from the mice stomach at Day 10 of scheme 1, 5 days after feeding *L. lactis* cloned with Laterosporulin and MM1-Laterosporulin, showed positive results when subjected to PCR with primers specific to the pTKR vector, proving that the bioengineered bacteria was present in the mice stomachs. This demonstrates that the bioengineered *L. lactis* remained for the duration of the therapy/study in the stomach with the plasmid still at a detectable amount (Figure 1)

**FIGURE 1.**
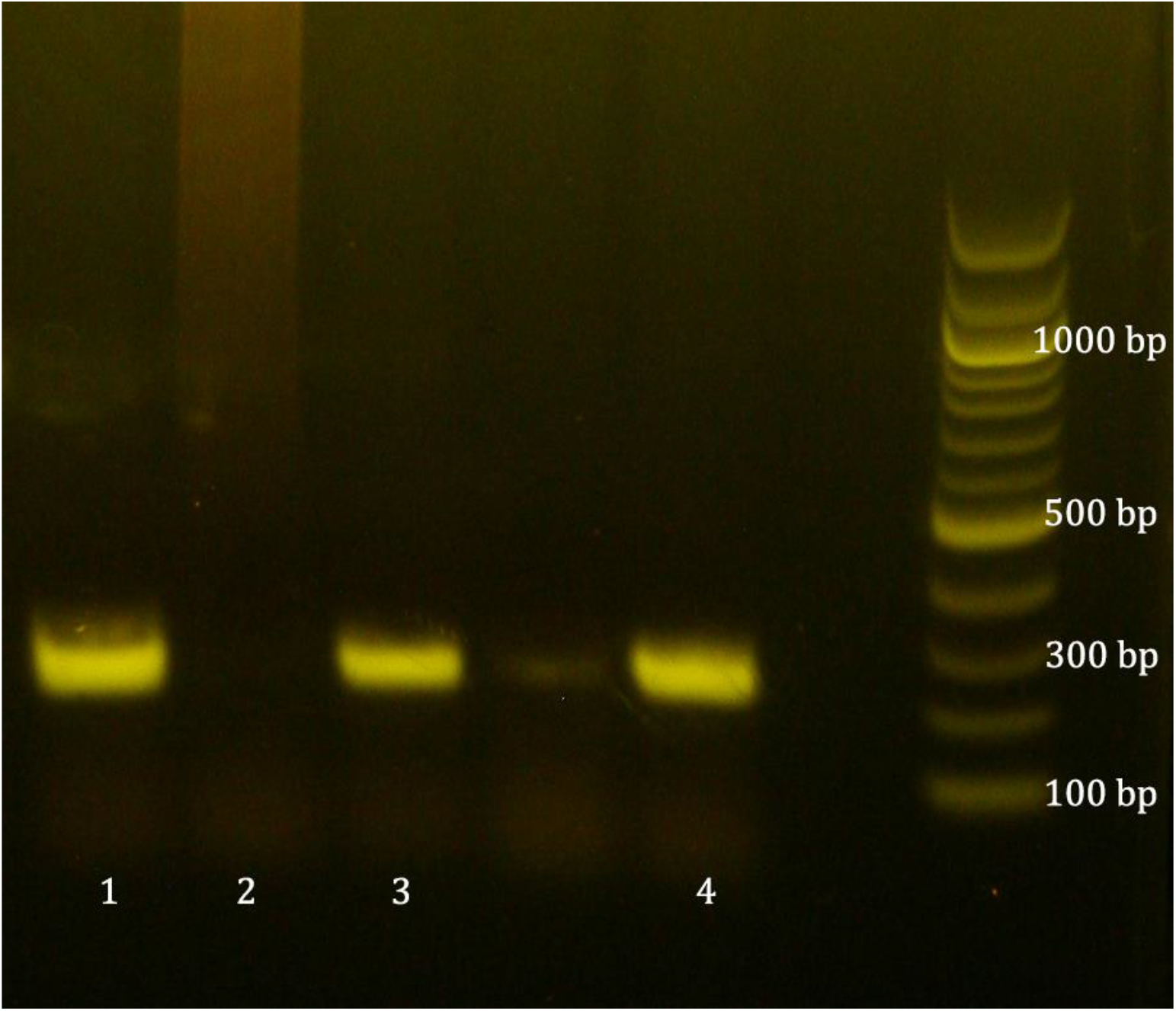
1% Agarose gel electrophoresis of stomach samples subjected to PCR with primers specific to the pTKR vector the *L. lactis* was cloned with. Lane 1) Positive control (200 ng of pTKR plasmid), 2) Negative control (mice stomach sample with no *L. lactis* inoculum, 3) Stomach sample at Day 10 from mice fed with Lat-*L. lactis*, 4) Stomach sample at Day 10 from mice fed with MM1-Lat-*L.lactis*

### Both gAMP and AMP releasing L. lactis reduced H. pylori load in mice stomach when used as a therapeutic

Mice stomach samples collected by reverse oral gavage method were subjected to qPCR to estimate the *H. pylori* load changing with time (Figure 2). The load reached its maxima at Day 5 after inoculation with *H. pylori* and had subsequent reduction after feeding *L. lactis* in the following 5 days except for the null control group and the group fed *L. lactis* with the empty pTKR vector. Both the gAMPs and AMPs releasing *L. lactis* had similar pattern of bactericidal activity with no significant difference between the AMP and their corresponding gAMP. At Day 10, the *H. pylori* titer of the extracted stomach sample were reduced by a factor 1/520 of the bacterial load in the empty vector group and 1/1100 of that of the null control group on average for all the 6 AMP and gAMPs. For the AMPs the titer value of *H. pylori* at Day 10 were on average reduced by a factor of 1/370 when compared to the empty vector group and by 1/800 when compared to the null control group on the same day. The gAMPs had a better reduction on average reducing *H. pylori* titer load by a factor of 1/860 against the empty vector group and by 1/1860 against the null control group for Day 10 samples. The titer of *H. pylori* for the gAMP-*L. lactis* were lower than the corresponding AMP-*L. lactis* but as stated, the values were not significantly different, echoing the findings of the *in vitro* assay in previous chapter.

**FIGURE 2.**
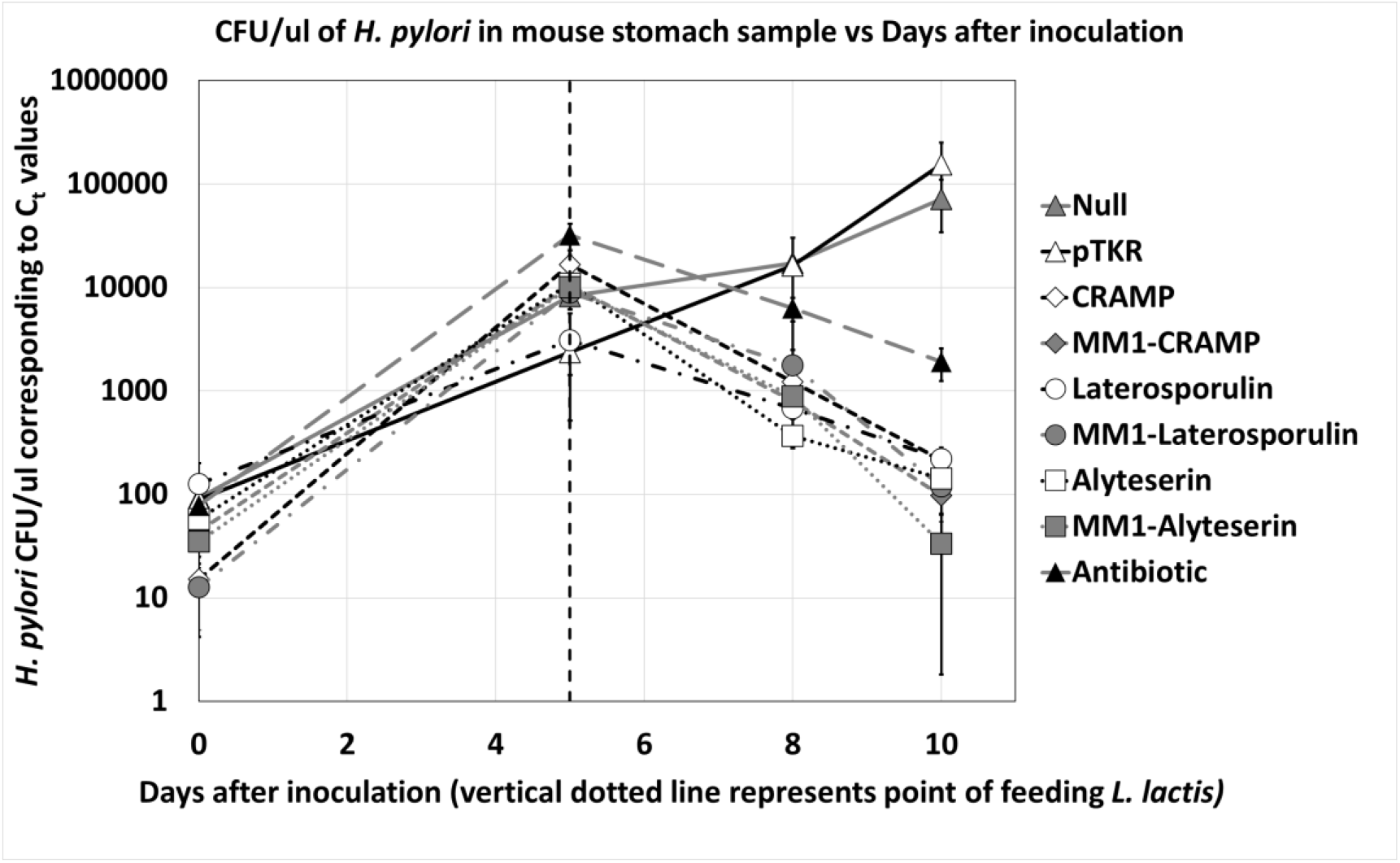
The CFU/μl of *H. pylori* found in the mice stomach, estimated from the corresponding C_t_ values determined by qPCR, plotted against days after inoculation with *H. pylori*. The dotted line at Day 5 represented the point of feeding *L. lactis* to the mice

### AMP and gAMP releasing L. lactis acts as a protective measure for the mice stomach

As we have seen with the therapeutic model, the AMP and gAMP cloned *L. lactis* helped reduce the load of *H. pylori* in the mice stomach compared to the null group and even the group fed with empty *L. lactis*. In the prophylactic approach, the observation wasn’t as drastic, but it did create a difference in the increase in *H. pylori* load in their stomach after feeding them with the bacterial inoculum for 3 consecutive days. As seen in Figure 3, the CFU/μl in the mice sample stomachs went up from Day 3, the day we began feeding the mice *H. pylori*, with the sharpest rise in the null group followed by the group being fed empty *L. lactis* as prophylactic. Both AMP and gAMP cloned *L. lactis* being fed as prophylactic 3 days before administering *H. pylori*, helped in reducing the ultimate titer of *H. pylori* in the stomach at Day 10 even though there was no significant discernible difference between the prophylactic effect between AMP and gAMP cloned *L. lactis*. On average both the AMPs and gAMPs when administered prophylactically had a lower yield of *H. pylori* in CFU/μl by a factor of 1/50 compared to the null group and 1/5 compared to the group fed empty *L. lactis*. Thus, we can conclude that the effect of the bioengineered *L. lactis* to reduce *H. pylori* load was more pronounced when used after the infection than before it happened as a prophylactic, but the approach still had some palliative effect.

**FIGURE 3.**
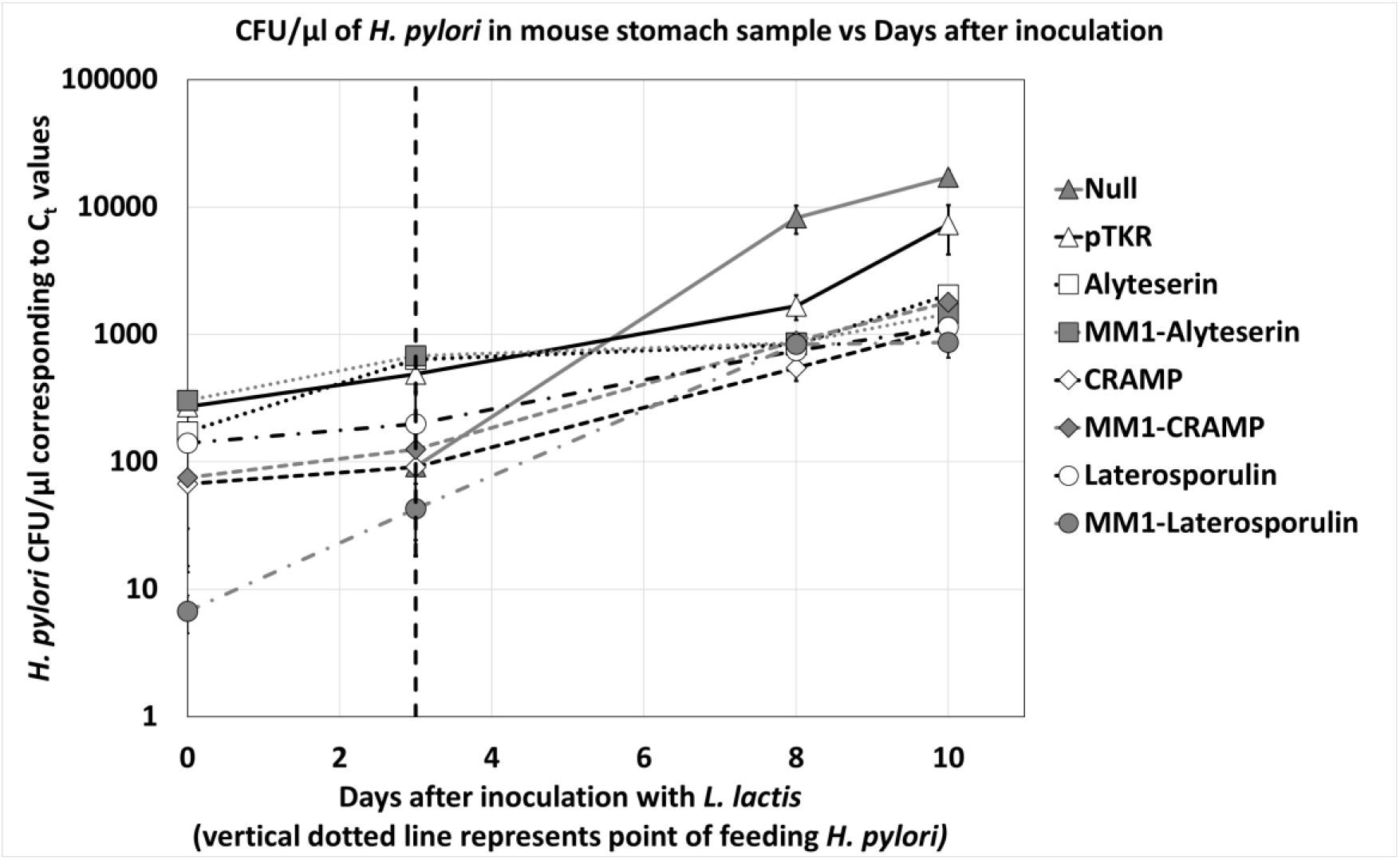
The CFU/μl of *H. pylori* found in the mice stomach, estimated from the corresponding C_t_ values determined by qPCR, plotted against days after inoculation with *H. pylori*. The dotted line at Day 3 represented the point of feeding *H. pylori* to the mice

### The gAMP cloned L. lactisreverses dysbiosis in mice stomach and restores microbiome diversity

Microbiome analysis through 16S rRNA gene sequencing of the 350 collected mice stomach samples revealed that *H. pylori* administration caused significant dysbiosis which was reversed by the subsequent feeding of cloned *L. lactis*. The dysbiosis was conspicuous on the samples taken 5 days after the beginning of administering H. pylori and was marked by a drastic drop in the diversity of species identified from the samples. In those samples a multitude of species/genera went down from over a few hundred to fewer than 10 across the subjects. The microbiome diversity was restored once feeding clone *L. lactis* of any variety-of AMP or gAMP, but subjects with gAMP-*L. lactis* had a significantly higher number of OTUs in all the temporal samples compared to AMP, empty vector, empty L. lactis and antibiotic treatment (Figure 4). Comparing the mean OTUs observed at sampling depth over 5000 sequences for each sample across the 10 day period, we observe that there was a marked decrease of the OTUs in all treatment past Day 5 and till Day 8 when the number of OTUs go back up for every treatment except for the Antibiotic and the AMP-*L. lactis* treatments and the Null Control group which received no treatment. As the microbiome diversity is restored, the mean OTUs observed in the sampled sequences revert to pre-*H. pylori* values for the rest of the treatment with the gAMP-*L. lactis* treated group showing the best and a significant recovery compared to the observed OTU values of the null control group, empty vector (pTKR) group, antibiotic group, or any of the AMP-*L. lactis* groups. Also, the observed OTU levels of AMP-*L. lactis* was actually not significantly different than the null control group results for the same day (Table 1), thus, proving that even if the AMP-*L. lactis* were successful in decreasing the *H. pylori* load, as seen in qPCR assays, they were not beneficial to the overall diversity of the stomach eco-system.

**FIGURE 4.**
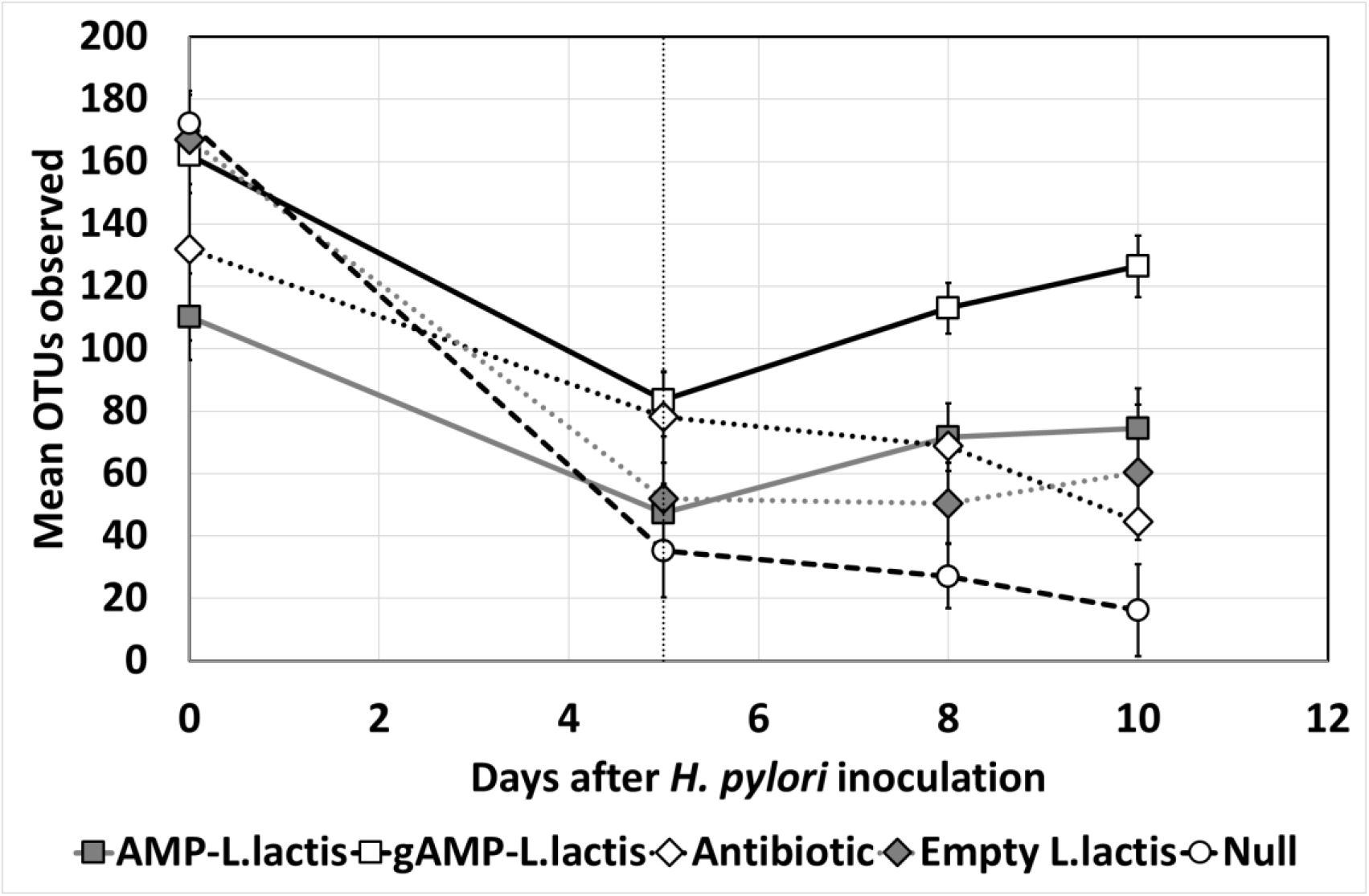
The mean Observed OTU numbers (at Genus level) for the therapeutic treatment groups. Each groups were fed *H. pylori* for 3 consecutive days beginning with Day 0 and they were fed their respective treatment (AMP-*L. lactis*, gAMP-*L. lactis*, Antibiotics, Empty vector-*L. lactis*) on Day 5 after sampling (denoted by the broken vertical line)

**TABLE 1.** The p-values of the t-test performed between observed OTU values (at Genus level) at day 10 for each treatment groups (AMP-*L. lactis*, gAMP-*L. lactis*, Antibiotics, Empty vector-*L. lactis* and Null Control)

| *p-values* | gAMP | AMP | Antibiotic | Empty |
| --- | --- | --- | --- | --- |
| Null | 4.04362E-45 | 7.75921E-29 | 4.88147E-20 | 2.9924E-12 |
| Empty | 0.000118242 | 0.296878437 | 4.98665E-40 |  |
| Antibiotic | 7.1317E-129 | 7.2617E-108 |  |  |
| AMP | 8.24156E-05 |  |  |  |

F or the prophylactic groups, all treatment groups including the Empty vector-*L.lactis* group had some protective effect in the mice stomach and the diversity with only the null control group having a plummeting effect in the OTU levels (Figure 5). Even here, the gAMP-*L.lactis* group had the best and recovery and the number of OTUs observed even surpassed the number of OTUs observed at Day 0 whereas the mean observed OTU levels pf AMP-*L. lactis* fell bellow that of Empty-*L.lactis* group.

**FIGURE 5.**
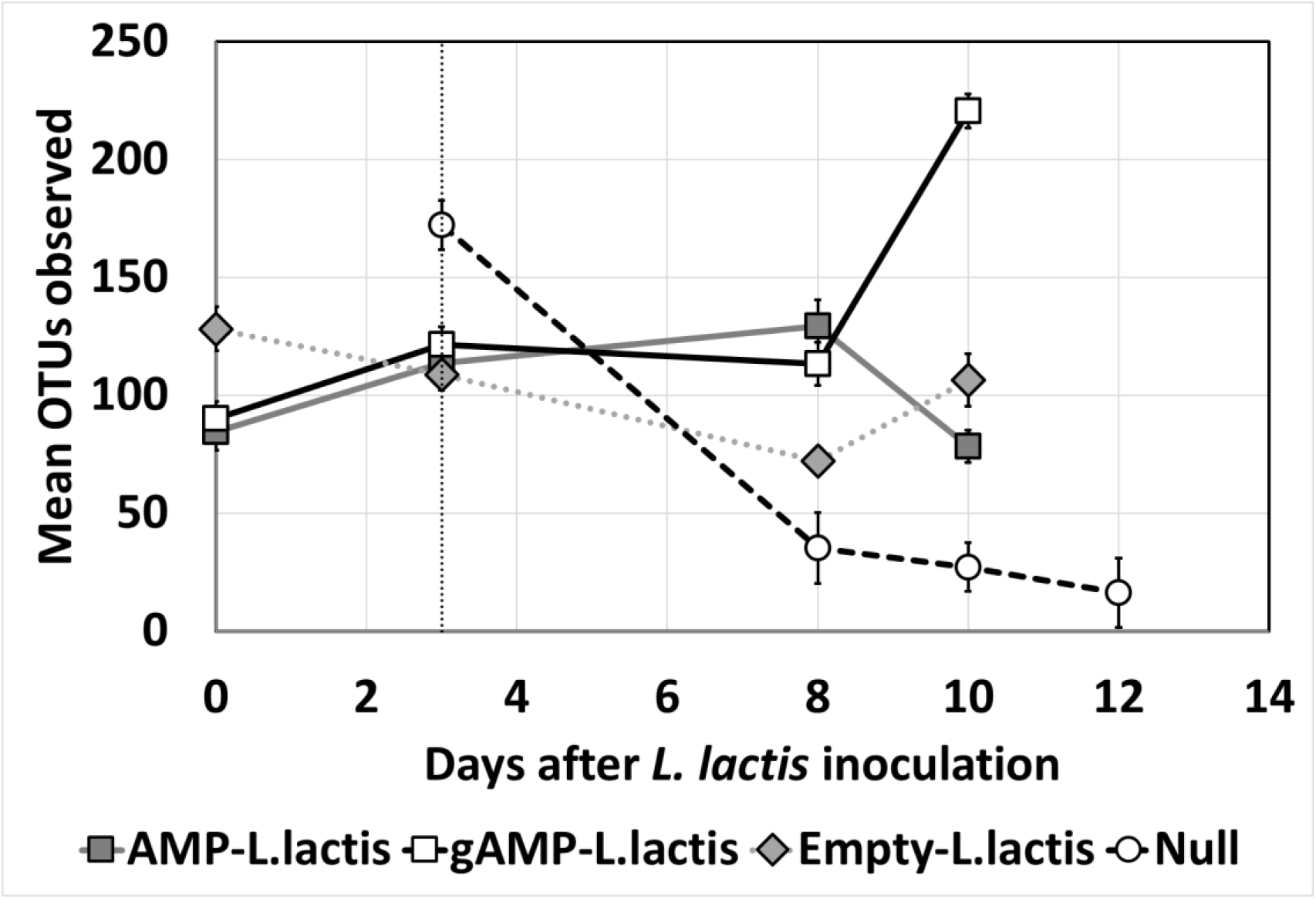
The mean Observed OTU numbers (at Genus level) for the prophylactic treatment groups. Each group were fed their respective treatment (AMP-*L. lactis*, gAMP-*L. lactis*, Antibiotics, Empty vector-*L. lactis*) on Day 0 and they were fed *H. pylori* for 3 consecutive days beginning with Day 3 after sampling (denoted by the broken vertical line)

### Microbial Dysbiois Index was created to examine the nature of recovery after H. pylori infected mice were fed bioengineered L. lactis

The relative abundance of all the sequenced taxa at Genus level were used to create a compositional analysis table with a subset of Day 0 samples and Day 5 samples, with the variable between them being the Day 5 samples were infected with *H. pylori*, which decimated the diversity of the mice stomach. The compositional data between the samples presented correlations between features (here, relative abundance of different taxa) due to the nonindependence of values that must sum to a fixed total. The CCREPE package was used to abrogate the correlation and determine the significance of a similarity measure for each feature pair using permutation/renormalization and bootstrapping (Faust et al., 2012; Schwager et al., n.d.). This generated an N-Dimensional checkerboard with similarity scores, p-values and false discovery rate q-values corrected for the effects of compositionality. The top features with best p-values and q-values (<0.12) were selected to create a correlation network with the features (Figure 6) based on their similarity scores (between 1 and −1 based on their correlation across the samples of Day 0 and Day 5). This revealed 8 genera being positively correlated with each other 2 other genera being negatively correlated with the other 8, based on their relative abundance across Day 0 samples and Day 5 samples. On analyzing the change in relative abundances of these 10 genera between the samples of Day 0 and Day 5, all the 8 positively correlated genera had a higher relative abundance in Day 0 compared to Day 5 and the other 2 had significantly higher abundance in Day 5 compared to Day 0 (Figure 7).

**FIGURE 6.**
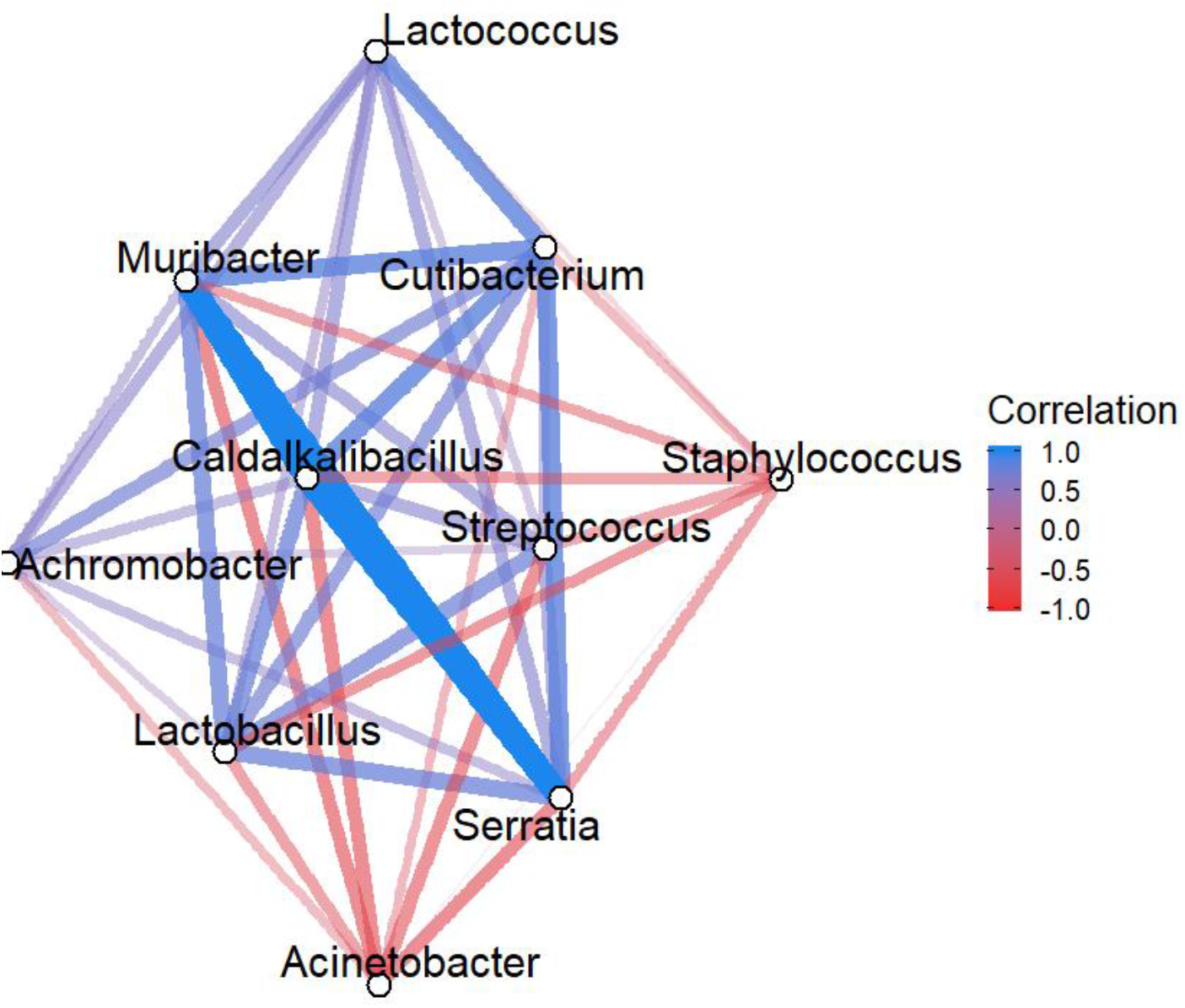
The 10 top genera showing significant similarity scores with each other across the Day 5 and Day 0 samples. The 8 genera (*Lactococcus, Muribacter, Cutibacterium, Caldalkaibacillus, Streptococcus, Achromobacter, Lactobacillus* and *Serratia*) had the most positive correlation with each other whereas *Staphylococcus* and *Acinetobacter* were negatively correlated with all the 8.

**FIGURE 7.**
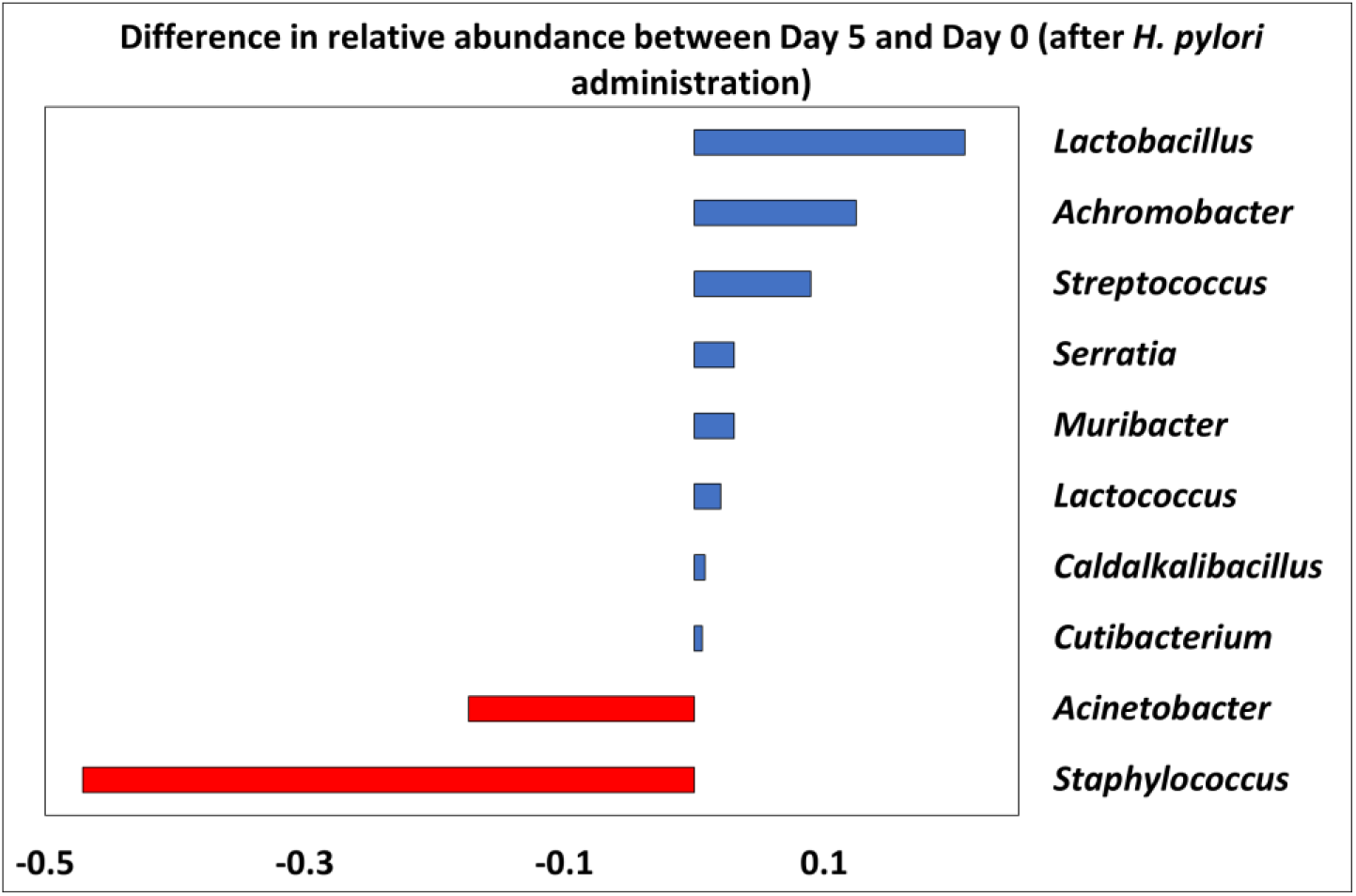
Difference in relative abundance of the 10 genera between Day 5 and Day 0 samples

The differential abundance of the 10 genera was used to create a Microbial Dysbiosis Index (MDI) using the formula:

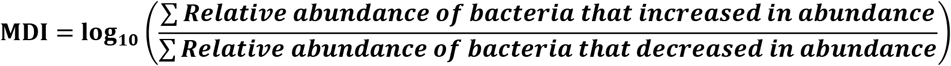

A positive MDI indicates a possible dysbiotic condition in which the 2 bacteria more abundant in the Day 5 (*H. pylori* infected) samples are overrepresented and the 8 bacteria that were positively correlated in the Day 0 samples, which resemble most the native microbiome, are underrepresented. Using the formula, the MDI for all the samples were calculated to observe the clustering of the samples based on the MDI and the relation it has with the log10 of the genera count for that sample (Figure 8)

**FIGURE 8.**
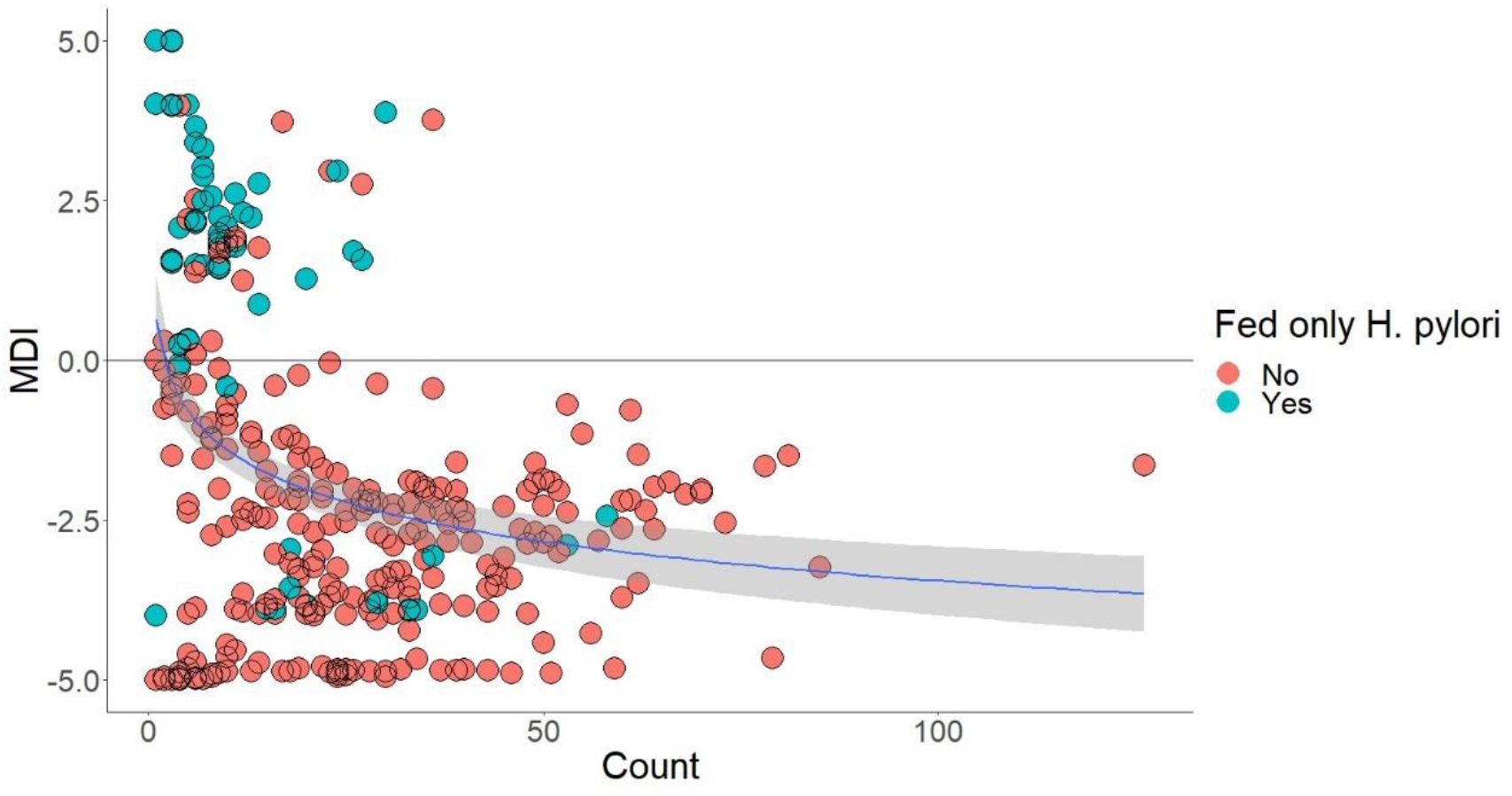
The MDI values of all samples plotted across the number of genera represented in the sample. The dots are colored representing whether the samples were from a mouse that have only been infected with *H. pylori* (Blue) or those who have had no bacteria fed or *L. lactis*/antibiotic fed after *H. pylori* (Red). The relation between log10 of genera count and is depicted by the blue line.

The model was validated by checking the prediction it made about the 349 samples (excluding the subset used for creating the model) on being dysbiotic (MDI>0) or non-dysbiotic (MDI>0), against whether the samples were really dysbiotic (infected with only *H. pylori*) or not. It correctly predicted 89.08% of samples based on their MDI.

The samples were then divided into Therapeutic (Figure 9) and Prophylactic (Figure 10) groups and checked whether which treatment affected the MDI scores/dysbiotic level of the mice that they were given to. For the Therapeutic group, most of the samples that have had only *H. pylori* being administered scored a positive MDI and the other most represented group in the positive MDI cluster were the samples from the mice that were given Empty vector-*L. lactis* in subsequent days after being infected with *H. pylori*. The *H. pylori* only fed samples with a negative MDI had significantly larger genera count that the rest of the similar *H. pylori* only samples.

**FIGURE 9.**
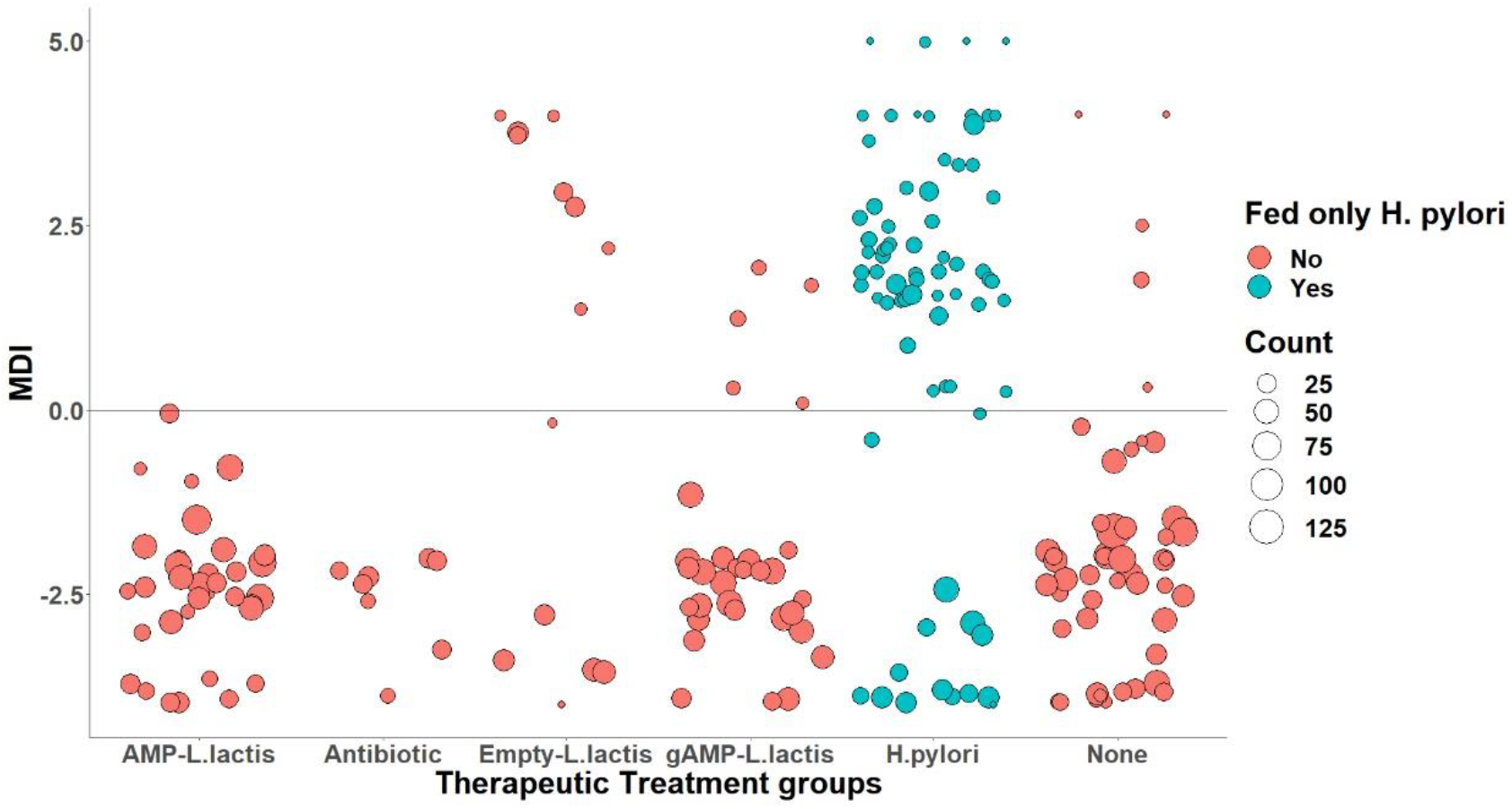
The samples from the Therapeutic group plotted according to their MDI scores. The dots are colored representing whether the samples were from a mouse that have only been infected with *H. pylori* (Blue) or those who have had no bacteria fed or *L. lactis/*antibiotic fed after *H. pylori* (Red). The size of the dots represents the number of genera represented in the samples.

**FIGURE 10.**
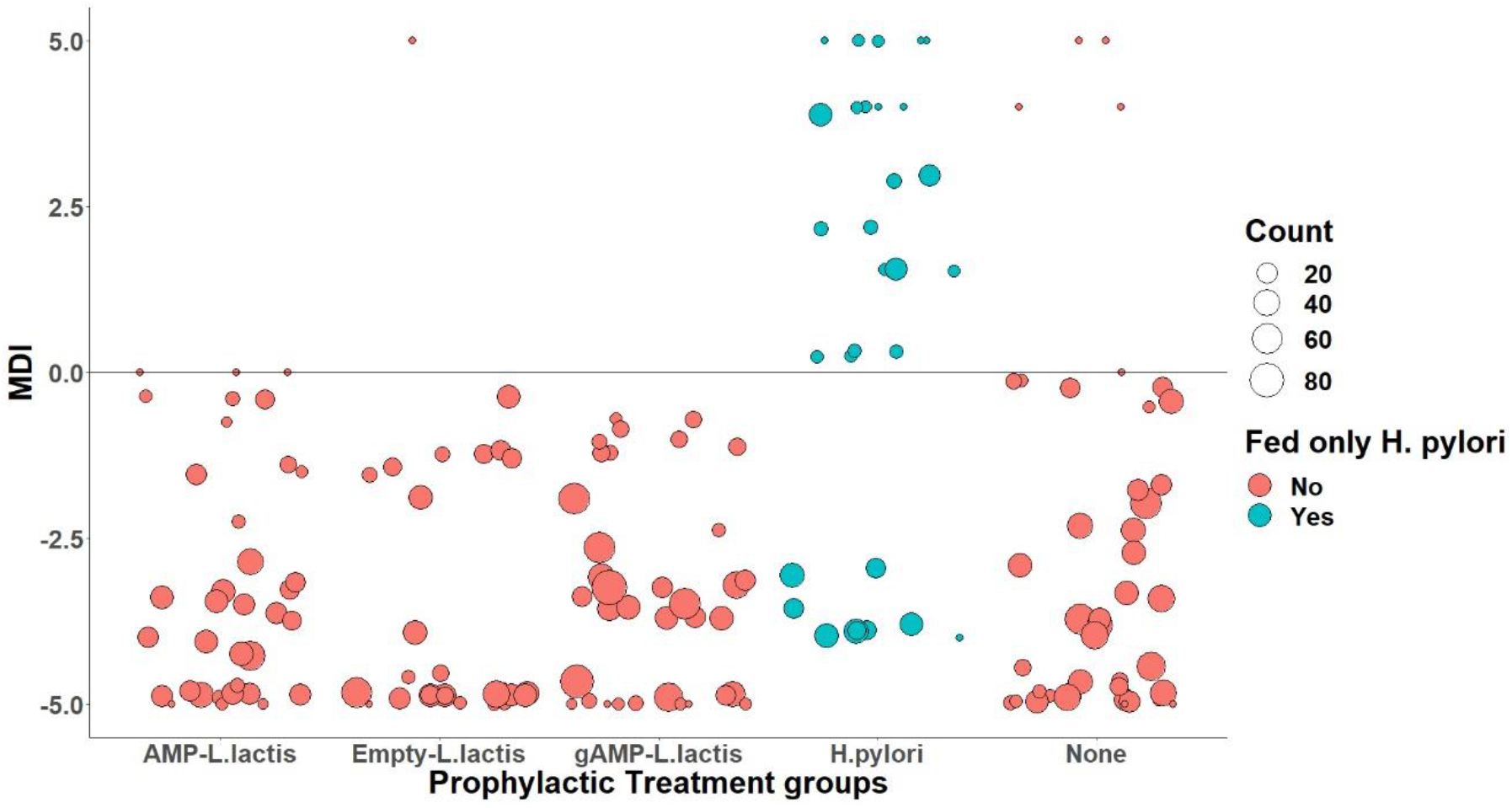
The samples from the Prophylactic group plotted according to their MDI scores. The dots are colored representing whether the samples were from a mouse that have only been infected with *H. pylori* (Blue) or those who have had no bacteria fed or *L. lactis/*antibiotic fed after *H. pylori* (Red). The size of the dots represents the number of genera represented in the samples

For the prophylactic groups, the MDI scores were negative for every sample belonging to mice that were primed with *L. lactis*, as seen in previous analyses, with the *H. pylori* only samples having mostly positive MDI score. These trends were also seen when the mean MDI scores of each treatment group in the Therapeutic and Prophylactic schemes were plotted against the Day the samples were taken. For the Therapeutic schemes (Figure 11), the MDI scores were negative for all treatments in Day 0, with all of them becoming positive on Day 5 after 3 consecutive days of administering *H. pylori*. On Day 8 and Day 10, for the AMP-*L. lactis*, gAMP-*L. lactis* and antibiotic treated groups, the scores returned to being negative while those of the Empty-*L.lactis* and null control group remained positive.

**FIGURE 11.**
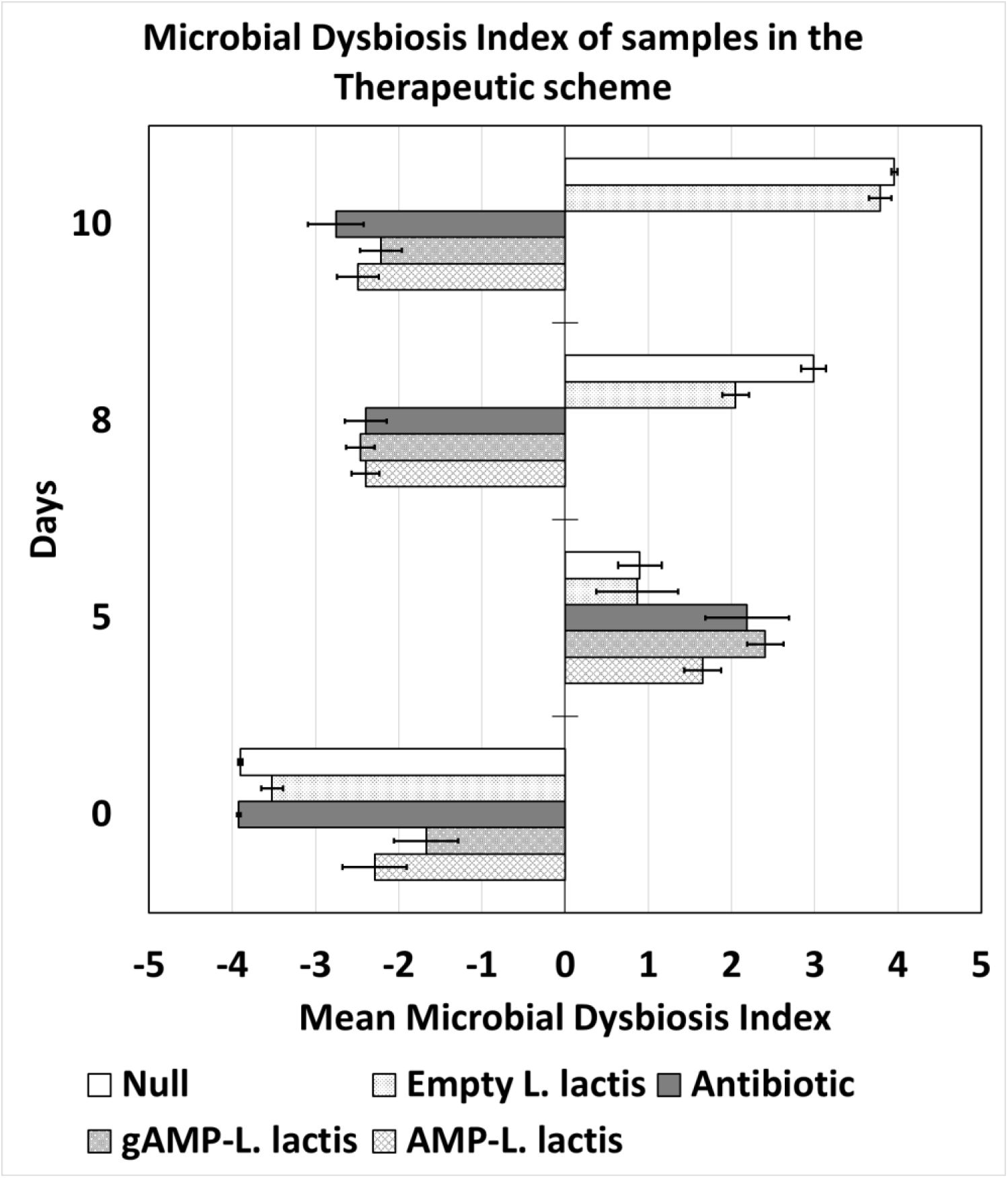
Mean MDI scores for each treatment group in the Therapeutic scheme against the day they were sampled.

**FIGURE 12.**
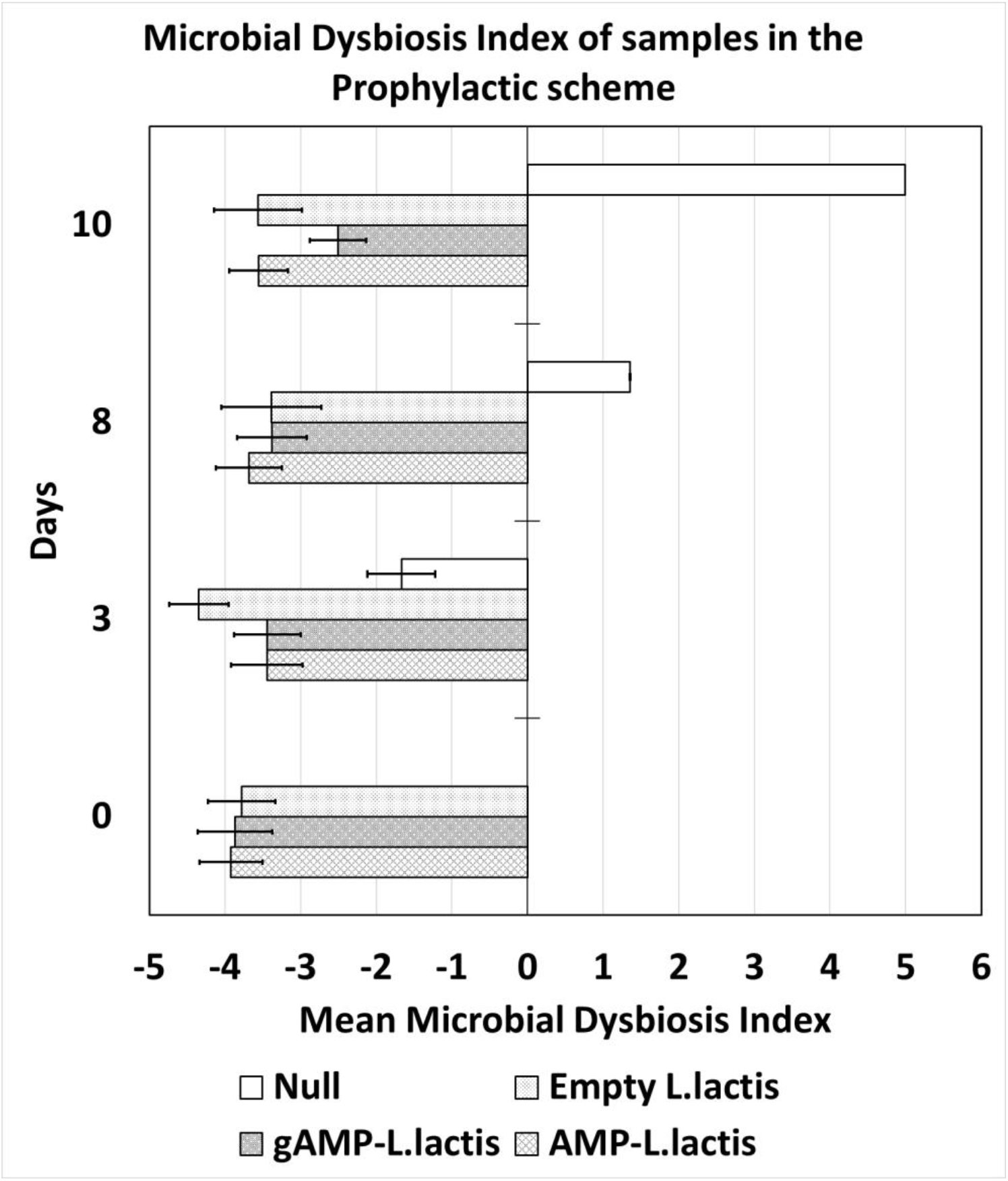
Mean MDI scores for each treatment group in the Prophylactic scheme against the day they were sampled

For the Prophylactic schemes, all the treatment groups had negative MDI scores at Day 0 and they remained negative at Day 3, 8 and 10, the AMP-*L. lactis*, gAMP-*L. lactis* and Empty-*L. lactis* groups. The null control group had negative MDI scores on Day 3, which turned positive on Day 8 and 10 after administering *H. pylori* for 3 consecutive days.

## Discussion

All the *in vivo* assays demonstrated that *L. lactis* cloned with AMP/gAMP were effective in killing *H. pylori*. The qPCR for the oral gavage samples from mice stomachs shows that both gAMP and AMP-*L. lactis* were similarly efficacious in reducing the *H. pylori* titre. This echoes what we saw through the *in vitro* co-culture studies in the previous chapter. The reduction was significant compared to the null control and even *L. lactis* cloned with empty pTKR vector which shows that the action of eliminating *H. pylori* is done by the AMP/gAMPs being released and not just by the presence of the *L. lactis* bacteria. The same effects were shown by the antibiotic treated group even thought the Day 10 CFU/μl of *H. pylori* in the samples were higher, not significantly, than either of the AMP or gAMP-*L. lactis* groups. For the prophylactic scheme, all the treatment groups including the Empty-*L. lactis* showed a significant protective effect on the mouse stomach microbiome when compared to the null control group, even though the *H. pylori* CFU/μl values at Day 10 were around 5 times on average of that of the CFU/μl values of the AMP/gAMP-*L. lactis* group. Thus, this shows that the bioengineered *L. lactis* can be used either as a therapeutic or prophylactic to recover or protect from an *H. pylori* infection in the mice stomachs with the therapeutic effect being more pronounced compared to the null group. In the therapeutic groups, the AMP/gAMP-*L. lactis* reduced the load of *H. pylori* significantly compared to the mice groups being fed Empty-*L. lactis*, hence proving that the bactericidal effect was brought upon by the release of the AMP or gAMPs from the bioengineered probiotic and not just the probiotic itself. As for the protective measure, the effect of the probiotic to prime the mice stomach microbiome for a dysbiotic shock like introduction of *H. pylori* was consistent despite the fact they were or not cloned with AMP/gAMPs. This might just be due to the fact of the innate bioprotective nature of *L. lactis* which might help in maintaining the pH of the mice stomach which is increased due to the colonization of *H. pylori*, and is essential for its survival, and also just out compete the introduced pathogen and maintain the native biodiversity.

Whether there was a difference in the impact of the AMP-*L. lactis* and the gAMP-*L. lactis* on the native micorbiota, this was determined by the 16s rRNA sequencing using Illumina MiSeq to see what changes they caused in the stomach microbiome. We could see that on Day 5 of our therapeutic scheme experiment, the mice that have been fed *H. pylori* for 3 days had a marked change in their microbiome diversity in which almost all the genera were wiped out except for *Staphylococcus* and *Acinetobacter*. According to prior literature, both these genera are linked to diseased condition in mice and hence the growth in their relative abundance is a major sign of dysbiosis (Zavros et al., 2002, Misawa et al., 2015). Even though the sequencing did not capture the corresponding rise in *H. pylori* level, mainly because the technique is not ideal for quantitative analysis and hence we used the qPCR, this remarkable drop in the abundance of other species may be induced by the conditions created by *H. pylori* infection like increasing stomach fluid pH, erosion of gastric mucosa which serves as a substrate for many gastric flora etc. *Staphylococcus* prefers alkaline environment to acidic and *Acinetobacter* has similar gastric colonization pattern as *H. pylori*, and often these two genera are found as concurrent bacterial flora in samples from patients with *H. pylori* induced hypochlorhydria, dyspepsia and gastritis. *Acinetobacter* causes gastritis and hypergastrinemia, like *H. pylori*, and their coexistence maybe due to their similar nature in sculpting gastric environment like increasing pH, creating inflammation and vacuolation of gastric tissue to enhance their survivability (Zavros et al., 2002). Thus, the rise of these two genera can be seen as an after effect of *H. pylori* colonization in the mice stomachs. These two genera were also represented in the top 10 genera represented in the 350 samples we sequenced from the mice stomach. Among them were *Lactobacillus, Streptococcus, Muribacter, Cutibacterium etc*, few genera often associated with healthy mice gastric and gut microbiome which often maintains mice gastric pH, lactate levels, metabolite homeostasis etc (Dargahi et al., 2020; Granland et al., 2020; Rocha Martin et al., 2019; Wang et al., 2018). Here we can see that the samples from the mice fed gAMP-*L. lactis* and AMP-*L. lactis* had a significantly better recovery of the relative abundance for these genera than empty *L. lactis*. The MDI equation predicted the dysbiotic (infected with only *H. pylori*) states of the mice stomach samples with 90% accuracy and hence could be used as a credible yardstick for the purpose. Plotting the MDI score distribution we saw that the therapeutic administration of AMP and gAMP-*L. lactis* had equivalent response in modulating the score and hence recovering from the dysbiosis.

On examining the diversity however, using the observed OTU number at genus level, we find that gAMP-*L. lactis* had the best recovery in the number of OTUs pre and post administration of *H. pylori*. This phenomenon was seen in both therapeutic and prophylactic schemes, with significant increase in observed OTU levels than AMP-*L. lactis*, Empty-*L. lactis* or antibiotic treatment at Day 10 of the regimen.It was also observed that the AMP-*L. lactis*, even though showing a recovered OTU level, was not significantly different than the effect of Empty-*L. lactis* at Day 10 for either the therapeutic or prophylactic schemes. The samples from the mice in the therapeutic groups that were fed gAMP-*L. lactis* consistently had the highest observed OTUs overall at almost all sampling depths. When we examine the number of OTUs observed beyond sampling depth of 5000, we see a day-wise trend in change of OTUs with everyone plummeting low till day 8, perhaps due to the *H. pylori* induced dysbiosis, and then a recovery in their numbers is only observed for the gAMP-*L. lactis*, empty *L. lactis* and null control. The downwards trend of the observed OTUs in samples from mice that were fed the antibiotic cocktail and the null group, with the AMP-*L. lactis* and Empty-*L. lactis* making a weaker recovery. This weaker effect of AMP-*L. lactis* was mirrored in the prophylactic group samples as well and even in Day 10 of the regimen, the mean number of OTUs even dropped below that of Empty-*L. lactis* fed sample. One possible reason behind this could be the fact that both the antibiotic treatment and the AMP released from the *L. lactis* indiscriminately kills the bacterial flora in the stomach causing an overall drop in the OTUs observed along with *H. pylori*, the intended target. But the gAMP-*L. lactis*, which also killed *H. pylori* as proved by the qPCR studies, had the highest observed OTU in Day 10, proving that they perhaps killed *H. pylori* with higher specificity over other bacteria in the stomach. The rise in number of OTUs in Empty *L. lactis* group on Day 10 may be due to the healing of microbiome on passage of time or the presence of a helpful probiotic species like *L. lactis* respectively. Thus, all the experiments and results we have examined in the chapters Three and Four help establish the fact that the *L. lactis* cloned with AMPs and gAMPs are effective in eliminating *H. pylori* both *in vitro* co-cultures and *in vivo* from mice stomachs infected with *H. pylori* but the gAMP releasing *L. lactis* eliminates *H. pylori* in a more selective manner as seen from its relative inaction against the non-target *E. coli* and *L. plantarum in vitro* and also having the highest overall number of OTUs observed in the mice stomach at the end of the experiment among all the treatments.

